# Penumbria: Advanced 3D cell segmentation for biomedical imaging

**DOI:** 10.64898/2026.06.30.735527

**Authors:** Laurids Stockert, Joseph Donovan, Herwig Baier

**Affiliations:** Max Planck Institute for Biological Intelligence, Genes – Circuits – Behavior, Am Klopferspitz 18, Martinsried, Germany

## Abstract

Quantitative analysis of three-dimensional cellular architecture is fundamental to understanding tissue organization, disease progression, and drug response. There are effective approaches for 2D segmentation, yet 3D cell segmentation remains a critical bottleneck due to diverse cell morphologies, low signal-to-noise ratios, and data scarcity. We introduce Penumbria, a general-purpose 3D cell segmentation framework that achieves state-of-the-art accuracy across morphologically distinct cell populations and imaging conditions in volumetric microscopy. Penumbria formulates segmentation as a regression problem on distances to cell boundaries, supporting instance reconstruction without shape priors and permitting end-to-end GPU inference. A U-Net-based architecture with xLSTM bottleneck blocks and patch embeddings enables multi-scale feature extraction, long-range modeling of spatial context, and convolutional feature-volume tokenization. The model is extended with two modules: a Global Zernike Phase Layer, which learns Zernike-parameterized phase corrections in the frequency domain to deal with optical aberrations such as defocus and tilt, and a Scaled Geocaps Layer, which samples features at fixed grid locations across multiple spatial scales, routing evidence between them such that a detection is only confident where concordance holds across scales simultaneously. Across four diverse 3D datasets selected to probe the limits of existing methods, Penumbria outperforms Cellpose-SAM on all four and StarDist-3D on three, with comparable accuracy on the fourth, achieving up to a 38% improvement in mean average precision over the second-best method. Penumbria’s strong boundary accuracy further supports downstream analyses such as quantifying membrane dynamics or protein localization. We also release a hand-labeled confocal dataset of 1,543 zebrafish neurons across seven volumes, on which Penumbria can be easily and quickly trained.

## Introduction

Object detection and segmentation can accelerate data throughput and reduce the error-prone, laborious task of manual image labeling, particularly for instance segmentation of cells from microscopy volumes. However, developing algorithms for 3D cell microscopy data is challenging due to factors such as low signal-to-noise ratio (SNR), scarcity of annotated data, high variability in cell shapes, axial anisotropy, algorithmic memory constraints, and the curse of dimensionality. Neural networks have helped to address some of these challenges through learned feature extraction and tailor-made training objectives [1, 2]. In conventional real-world image tasks, a popular approach is to pre-train on large image datasets to learn relevant, reusable features [3, 4]. However, this strategy is less feasible in biomedical imaging due to substantial shifts in data domains across microscopy modalities [5]. Therefore, researchers often need to train custom models from scratch on their own datasets, which has led to the development of many specialized architectures that tackle cell segmentation from different structural viewpoints. These broadly fall into two families: proposal-based and proposal-free.

In proposal-based pipelines, candidate regions are identified first, and masks are refined from those proposals, as in R-CNN and Fast R-CNN [6, 7]. Focal loss [8] later addressed the persistent class imbalance between background and foreground pixels, helping drive the development of single-pass detectors such as YOLO [9]. Yet accurate segmentation depends on accurate proposals, and errors introduced at the proposal stage cannot be corrected downstream. This chicken-and-egg problem, combined with the sequential evaluation of bounding boxes, limits both parallelization and training efficiency [10, 11].

Proposal-free methods sidestep these issues and now dominate the field. CellPose, for example, predicts flow fields from a heat-diffusion simulation and integrates them to recover cell shapes [1]. EmbedSeg instead learns pixel embeddings that cluster by object identity, with offset vectors and anisotropic bandwidths to handle differences in cell size [12, 13]. Watershed-based methods remain a practical choice for crowded scenes and are routinely applied as post-processing after heatmap regression [14, 15, 16, 17, 18, 19, 20]. StarDist-3D represents cells as star-convex polyhedra, densely predicting boundary distances along rays evenly distributed over a reference ellipsoid fitted to the data, with final shapes obtained via non-maximum suppression [21, 22]. This works well when convexity holds, but less so when it does not.

Each of these methods navigates specific trade-offs: CellPose adapts to varied morphologies but expands predictions locally from a single point, whereas StarDist-3D imposes a convex prototypical shape prior beneficial for analyzing low-contrast imaging that simultaneously limits shape flexibility. Recently, CellPose has integrated SAM [23, 24, 25] to leverage large-scale pretraining. Even so, aggregating 2D features for volumetric data remains no substitute for natively 3D expressive power.

Here, we present Penumbria, a cell segmentation pipeline optimized for three-dimensional microscopy that achieves high morphological fidelity spanning varied shapes without compromising robustness under challenging imaging conditions. Across four diverse 3D datasets, each difficult enough to avoid saturation yet capable of exposing specific contextual weaknesses, we consistently outperform Cellpose-SAM and match or exceed StarDist-3D, achieving up to a 38% gain in mean average precision while training entirely from scratch.

## Results

### Design choices for Penumbria

Penumbria is built around two design principles: shape flexibility and adaptation to different imaging conditions. For shape flexibility, we employ a classic combination of heatmap regression followed by seeded watershed flooding [14]. Continuous image regression represents a smooth optimization target for neural networks while allowing greyscale morphological operations to facilitate iterative region growing under local connectivity constraints. Conceptually, this can be imagined as framing the task as a topological problem, where local extrema signify the centers of individual objects that are flooded in a controlled manner to establish boundaries. We draw on methods developed by Luc Vincent, using the h-maxima transform for seed detection prior to watershed segmentation [26, 27]. In particular, the h-maxima transform highlights local extrema that are not only significant but also well-connected, conferring robustness to internal brightness variabilities within cells and guarding against erroneous splitting.

Complementing this algorithmic choice is a neural network that remains highly competitive under various signal perturbations: U-vixLSTM [28]. It represents the product of focused research into expressive sequence modeling using recurrent networks and a well-established multi-scale feature extraction via a Unet architecture [29, 30, 31]. This is part of a broader pattern currently emerging in computer vision, as we see models based on hybrid architectures, such as Segmamba or Segformer, increasingly outperform single-paradigm methods [32, 33]. This is no coincidence, as CNNs and sequential networks tackle different aspects of visual feature extraction, such as texture identification or global coherence, respectively [34, 35, 36].

Building on U-vixLSTM, we introduce two novel architectural contributions alongside a key modification. First, we replace BatchNorm with Filter Response Normalization (FRN) [37, 38]. The former needs larger batch sizes to yield stable estimates of mean and variance across features, which is infeasible for true three-dimensional processing. FRN, by contrast, can normalize feature scales while keeping brightness cues intact and avoiding bias drift via threshold linear units.

Secondly, we develop a Global Zernike Layer situated at the input stage to account for optical variability across different microscopes. This layer corrects optical aberrations such as tilt or defocus by applying a phase correction directly in the frequency domain, acting as a physical pre-processor that standardizes volumes before they enter the main segmentation backbone. Its mathematical basis lies in the Zernike polynomials, which form an orthogonal basis on the unit disc (i.e., the microscope lens) [39, 40]. Traditionally defined in 2D, these polynomials can be extended to 3D [41].

Finally, we propose a Scaled Grid Geocaps layer which samples features at fixed grid locations across multiple spatial scales, routing evidence between them such that a detection is only confident where concordance holds across scales simultaneously. This borrows the routing mechanism from Hinton’s Capsule Networks [42], but transfers it from a semantic context, where capsules establish agreement between object-part relationships, to a geometric one, where agreement is sought between the same spatial location observed at different scales.

Crucially, this layer is directly fed to the xLSTM bottleneck alongside the longer Unet encoder path, exploiting the module’s high spatial expressivity. To stabilize the induced latent representation, we regularize the xLSTM input via localized feature distillation, which is loosely similar to teacher-student feature map distillation proposed by [43], except it provides individual shape hints. Concretely, this entails occasionally choosing a random cell during a training pass and relaying an “imprint” of its heatmap label directly to the xLSTM. Analogically, this can be thought of as a visual “afterimage” one sees after staring at an object for a long time and closing one’s eyes, driving the xLSTM to organize its representations in a spatially meaningful and coherent way, since the injected label is itself spatially coherent. An overview of the complete architecture is shown in Figure 1.

**Figure 1:**
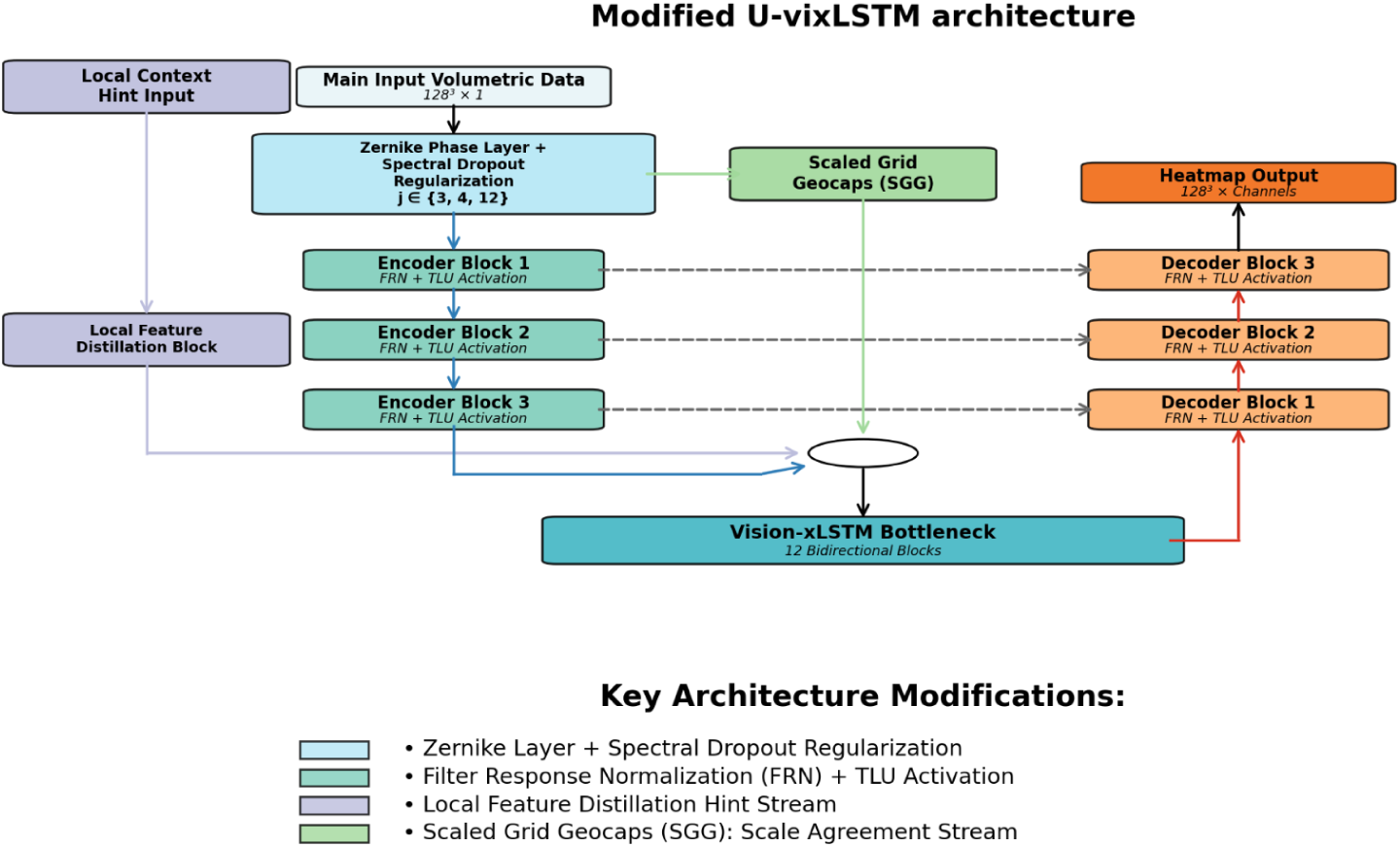
Illustration of the modified U-vixLSTM architecture with Zernike phase layer and Scaled Grid Geocaps (SGG). The input, an image cube with a sidelength of 128 and 1 color channel in this example, is first passed to the Zernike phase layer that includes spectral dropout, followed by successive downsampling and Filter Response Normalization with threshold linear units as activation functions. At the bottleneck, the xLSTM processes the heavily compressed feature volume sequentially and bidirectionally, before passing it to the decoder. Here, the SGG layer also introduces recursively aligned multi-scale features sampled at fixed frequencies, which are regularized by a local feature distillation block that occasionally provides local hints for cells, making the representations entering the xLSTM more spatially coherent. Skip connections maintain information flow between the encoder and decoder.

**Figure 2:**
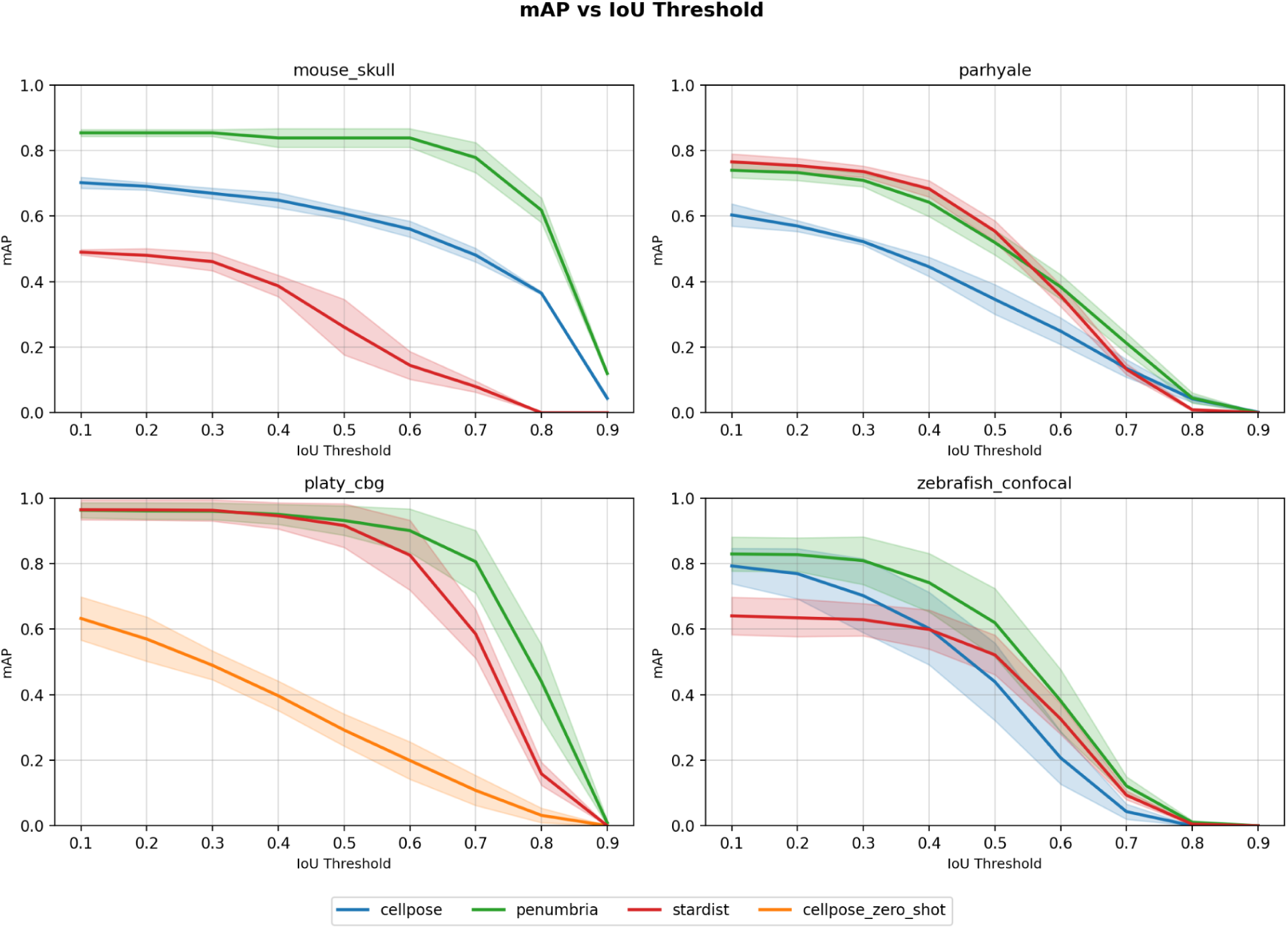
Mean average precision as a function of IoU threshold for Penumbria, StarDist-3D, Cellpose-SAM, and zero-shot Cellpose-SAM across all four benchmark datasets. Shaded regions indicate one standard deviation across images.

### Evaluation on challenging 3D microscopy datasets

We conducted baseline comparisons on four 3D datasets selected to capture the diversity of challenges in 3D segmentation. The datasets cover multiple imaging modalities, cell types, and densities, and include cells with eccentric shapes, anisotropic resolution, and intracellular features such as nuclei, chromatin, and invaginations that can resemble cell boundaries. In addition, background textures and imaging artifacts may mimic foreground structures, providing realistic obstacles that reveal performance differences and algorithmic limitations. Among these datasets, two were provided by [12]: Platynereis-Nuclei-CBG and Mouse-Skull-Nuclei-CBG. Furthermore, we benchmarked our model on the Parhyale-Nuclei-IGFL dataset [44] and our own dataset of confocal Zebrafish neurons. For an overview of the datasets, refer to Table 1.

**Table 1:** Organism and cell types, imaging modalities, and key challenges of datasets used for benchmarking.

| Dataset | Organism | Cell Type | Imaging Modality | Key Challenges | Source |
| --- | --- | --- | --- | --- | --- |
| Platynereis-Nuclei-CBG | *Platynereis dumerilii* | Nuclei | Light-sheet (LSM) | Dense packing, background texture mimicking foreground, high cell size variability, blob-like brighter imaging artifacts | [12] |
| Mouse-Skull-Nuclei-CBG | *Mus musculus* | Nuclei | Confocal | Extremely eccentric cell shapes exceeding network receptive field, internal nuclei structures (bright and dark) | [12] |
| Parhyale-Nucl<br>ei-IGFL | *Parhyale hawaiiensis* | Nuclei | Confocal | Low contrast, dark internal structures, very high anisotropy | [44] |
| Confocal<br>Zebrafish | *Danio rerio* | Neuronal nuclei | Confocal | High anisotropy, extremely dense cell packing | This work |

We evaluated each algorithm on the same data folds across runs, and performed three runs per algorithm-dataset pair. Performance was measured using mean average precision (MaP) at intersection-over-union (IoU) thresholds between 0.1 and 0.9, and the standard deviation was calculated for each threshold across all images. MaP is calculated by dividing the number of true positives by the sum of true positives, false positives, and false negatives. Moreover, we calculated the percentage-wise lead of the best-performing method on each dataset by averaging MaP values across all IoU thresholds. Finally, we computed the high-to-low (H/L) ratio, a measure of performance consistency obtained by summing the MaPs in the higher range (0.6-0.9) and dividing them by the sum of the MaPs in the lower range (0.1-0.4).

### Consistent performance gains across datasets and IoU thresholds

To evaluate Penumbria, we compared it against two established approaches: CellPose-SAM [24] and StarDist-3D [22]. The baseline methods were trained with their recommended settings (supplementary Material). Penumbria achieves higher performance metrics than both Cellpose-SAM and StarDist-3D across most datasets (see Table 2). Furthermore, it demonstrates a superior H/L ratio, indicating higher performance consistency across IoU thresholds. For Mouse Skull, Penumbria achieves an average MaP of 0.732, a 38.1% improvement over Cellpose-SAM, with minimal decline across the 0.1-0.6 IoU range. StarDist-3D records a lower average mAP of 0.256 in comparison.

**Table 2:** Mean average precision (TP / (TP+FP+FN)) over IoU-thresholds 0.1 - 0.9 for Penumbria, StarDist-3D, and Cellpose-SAM. The percentage lead denotes the average percentage difference between the best and second-best methods, computed by averaging MaPs across all IoU thresholds. H/L indicates consistency over all IoU-thresholds (higher -> better).

|  | 0.1 | 0.2 | 0.3 | 0.4 | 0.5 | 0.6 | 0.7 | 0.8 | 0.9 | Mean | % Lead | H/L Ratio |
| --- | --- | --- | --- | --- | --- | --- | --- | --- | --- | --- | --- | --- |
| **MOUSE_SKULL** |  |  |  |  |  |  |  |  |  |  |  |  |
| cellpose | 0.702<br>(±0.017) | 0.690<br>(±0.011) | 0.669<br>(±0.016) | 0.648<br>(±0.023) | 0.608<br>(±0.018) | 0.560<br>(±0.024) | 0.481<br>(±0.021) | 0.365<br>(±0.004) | 0.044<br>(±0.006) | 0.530 | — | 0.535 |
| penumbria | **0.854**<br>(±0.010) | **0.854**<br>(±0.010) | **0.854**<br>(±0.010) | **0.838**<br>(±0.028) | **0.838**<br>(±0.028) | **0.838**<br>(±0.028) | **0.778**<br>(±0.046) | **0.618**<br>(±0.038) | **0.120**<br>(±0.007) | **0.732** | **+38.1%** | **0.693** |
| stardist | 0.490<br>(±0.008) | 0.480<br>(±0.021) | 0.461<br>(±0.028) | 0.387<br>(±0.033) | 0.261<br>(±0.085) | 0.145<br>(±0.042) | 0.080<br>(±0.017) | 0.000<br>(±0.000) | 0.000<br>(±0.000) | 0.256 | — | 0.123 |
| **PARHYALE** |  |  |  |  |  |  |  |  |  |  |  |  |
| cellpose | 0.603<br>(±0.034) | 0.569<br>(±0.017) | 0.522<br>(±0.010) | 0.445<br>(±0.029) | 0.346<br>(±0.045) | 0.249<br>(±0.041) | 0.135<br>(±0.027) | 0.042<br>(±0.010) | **0.002**<br>(±0.002) | 0.324 | — | 0.200 |
| penumbria | 0.740<br>(±0.022) | 0.732<br>(±0.024) | 0.709<br>(±0.020) | 0.642<br>(±0.044) | 0.519<br>(±0.038) | **0.384**<br>(±0.037) | **0.212**<br>(±0.030) | **0.045**<br>(±0.016) | 0.000<br>(±0.000) | **0.443** | **Tie** | **0.227** |
| stardist | **0.765**<br>(±0.024) | **0.754**<br>(±0.022) | **0.736**<br>(±0.017) | **0.683**<br>(±0.025) | **0.554**<br>(±0.032) | 0.357<br>(±0.033) | 0.133<br>(±0.017) | 0.009<br>(±0.004) | 0.000<br>(±0.000) | **0.443** | **Tie** | 0.170 |
| **PLATY_CBG** |  |  |  |  |  |  |  |  |  |  |  |  |
| cellpose_zero_shot | 0.632<br>(±0.066) | 0.570<br>(±0.068) | 0.489<br>(±0.044) | 0.396<br>(±0.045) | 0.292<br>(±0.049) | 0.199<br>(±0.057) | 0.108<br>(±0.046) | 0.032<br>(±0.022) | 0.000<br>(±0.001) | 0.302 | — | 0.162 |
| penumbria | 0.963<br>(±0.023) | 0.960<br>(±0.025) | 0.960<br>(±0.025) | **0.950**<br>(±0.030) | **0.931**<br>(±0.045) | **0.900**<br>(±0.066) | **0.806**<br>(±0.095) | **0.440**<br>(±0.112) | **0.009**<br>(±0.008) | **0.769** | **+9.5%** | **0.562** |
| stardist | **0.964**<br>(±0.030) | **0.964**<br>(±0.031) | **0.962**<br>(±0.033) | 0.946<br>(±0.040) | 0.916<br>(±0.067) | 0.826<br>(±0.107) | 0.585<br>(±0.074) | 0.158<br>(±0.035) | 0.000<br>(±0.000) | 0.702 | — | 0.409 |
| **ZEBRAFISH_CONFOCAL** |  |  |  |  |  |  |  |  |  |  |  |  |
| cellpose | 0.793<br>(±0.054) | 0.769<br>(±0.076) | 0.702<br>(±0.113) | 0.602<br>(±0.111) | 0.439<br>(±0.118) | 0.207<br>(±0.080) | 0.043<br>(±0.023) | 0.000<br>(±0.000) | **0.000**<br>(±0.000) | 0.395 | — | 0.087 |
| penumbria | **0.829**<br>(±0.052) | **0.827**<br>(±0.051) | **0.809**<br>(±0.073) | **0.742**<br>(±0.089) | **0.619**<br>(±0.105) | **0.381**<br>(±0.095) | **0.122**<br>(±0.028) | **0.011**<br>(±0.005) | **0.000**<br>(±0.000) | **0.482** | **+22.0%** | 0.160 |
| stardist | 0.640<br>(±0.057) | 0.634<br>(±0.057) | 0.629<br>(±0.049) | 0.599<br>(±0.060) | 0.521<br>(±0.060) | 0.326<br>(±0.047) | 0.093<br>(±0.014) | 0.005<br>(±0.002) | **0.000**<br>(±0.000) | 0.383 | — | **0.169** |

On Parhyale, Penumbria ties StarDist-3D at a mean mAP of 0.443, with complementary profiles: StarDist-3D leads at lower IoU thresholds while Penumbria maintains performance at stricter overlap requirements, which require pixel-accurate boundaries to retain detections. To our knowledge, this is the first method to match StarDist-3D on Parhyale, a dataset whose relatively regular nuclear shapes align well with StarDist-3D’s star-convex shape prior. Cellpose records lower results here with an average MaP of 0.324.

For the Platy-CBG dataset, Penumbria and StarDist-3D show similar trends across lower IoU thresholds. Still, Penumbria maintains higher scores at stringent overlap requirements, resulting in an average mAP gain of 9.5%. Cellpose-SAM produced much lower results here when finetuned, with the model latching onto background texture and generating many false positives across multiple configurations. We replicated this failure on multiple machines. The behavior appears connected to the SAM-based release rather than the Cellpose approach generally. For context, [12] benchmarked an earlier Cellpose version on this dataset without the texture-latching failure. We therefore report Cellpose-SAM zero-shot performance. Finally, for the Zebrafish neuron confocal dataset, Penumbria maintains an average lead of 22% over CellPose. Between CellPose and StarDist-3D, the former exhibits higher scores at lower IoU thresholds, while the latter shows higher values at higher IoU thresholds.

## Discussion

This study aimed to develop and evaluate Penumbria, a 3D segmentation pipeline designed to address challenges arising from varying SNR and irregular cell shapes. By framing the segmentation task as a regression of re-scaled Euclidean distance transforms (EDT), we avoided many of the discretization issues found in standard segmentation methods. The U-vixLSTM architecture is central to this approach; placing xLSTM blocks at the bottleneck enables the model to capture the global context needed to filter out false positives effectively [28]. Moreover, we introduced one key modification and two architectural innovations.

First, we showed that the inclusion of a Global Zernike Layer at the input stage provides a learnable correction for optical aberrations, as an ablation analysis demonstrated small but consistent performance gains across different imaging conditions (Online Methods).

Interestingly, this happens for a fixed set of Zernike moments with j∈{3,4,12}. These low-order, primarily rotationally symmetric modes represent consistent distortions, such as defocus and spherical aberration. In contrast, higher-order polynomials tend to pick up on stochastic noise; for example, including j=24 can introduce artificial ringing artifacts in the background.

Secondly, we could extend the concept of capsule networks [42] from Gestalt-like perceptual hierarchy concordance to a geometric context, where recursively aligning coarse-to-fine grid structures on regularly sampled frequencies improves spatial coherence when regularized by local feature distillation in the form of occasionally injected label hints. Further ablation experiments demonstrated that this layer is particularly important for preserving coherent shapes of large objects beyond the network’s effective receptive field (Online Methods).

Finally, we also found that replacing BatchNorm with Filter Response Normalization (FRN) was a practical necessity for 3D work. FRN remains stable during high-memory 3D training with small batch sizes and preserves the brightness cues that EDT regression relies on more effectively than other normalization methods [38].

Reflecting these design choices, our benchmarks showed that Penumbria consistently surpassed Cellpose-SAM and either matched or outperformed StarDist-3D, with MaP gains of 9%-38% across most datasets. The data reveals two specific strengths. First, Penumbria handles initial cell detection well, as seen in the high scores at lower IoU thresholds. Second, it maintains a higher accuracy even as overlap requirements become more stringent (IoU ≥ 0.6). This makes the tool versatile for both simple counting tasks and more demanding work, such as membrane dynamics and protein localization, where the exact shape of the cell boundary is the primary focus. Finally, it is worth noting that Penumbria’s effectiveness depends on the availability of training data, as the morphological postprocessing parameters require tuning.

We observe that these performance differences are tied to the underlying architectures of each method. Cellpose, for example, relies on a flow-gradient generation strategy originally developed for 2D. While this allows the model to leverage extensive 2D pre-training on diverse datasets, the subsequent integration of these 2D predictions into a 3D volume is fundamentally limited by a lack of axial coherence. Additionally, while 2D Cellpose can use Vision Transformers (ViT), the cubic scaling of sequence length in 3D makes a natively higher-dimensional extension computationally infeasible for now.

Similarly, StarDist-3D has its own set of constraints. Using radial distance regression to incorporate shape priors is a clever strategy, but it struggles with the curse of dimensionality in 3D. Regressing 97 parameters per voxel requires downsampling the image by a factor of 4 to fit the model into GPU memory. This loss of resolution explains why StarDist-3D scores tend to drop off at higher IoU thresholds, where precision is key. This effect becomes even more pronounced when cells have eccentric or non-convex shapes, as the star-convex assumption no longer holds.

By contrast, Penumbria demonstrates that a natively 3D approach with minimal shape priors and morphologically adaptive postprocessing offers a clear advantage in handling complex microscopy volumes, providing researchers with a versatile tool for effective 3D segmentation in dense samples. It is thus particularly relevant in areas of biomedicine where training data are available, and high morphological fidelity is critical.

## Materials and Methods

Our method is most similar to the work carried out by Scherr et al [20] and Lux and Matula [14]. Both formulate cell segmentation as a regression task but differ in their regression targets, post-processing, and applicability, and are discussed in turn. Scherr and colleagues developed a general-purpose segmentation method for 2D and 3D that is highly performant in the competitive cell tracking challenge, lending credence to its versatility. They use a UNet with two decoders, where each decoder learns a specialized heatmap: one for object shapes generated by a cell-wise Euclidean distance transform, and another for Euclidean distance to neighboring cells. Once these targets are learned, instance labels are obtained by seeding the image with a thresholded version of itself and then applying watershed flooding. However, the contribution of the neighbor distance map to segmentation performance has been evaluated only partially, and evidence that it improves results across different cell types and imaging conditions is lacking.

Lux and Matula, by comparison, exclusively work on 2D images and employ a simpler target learned by two CNNs: one learning to distinguish foreground from background, and the other to predict watershed markers, which can be conceptualized as the foreground minus a uniformly thick boundary. To obtain the seeds for watershed flooding, however, they use a classical operation called the morphological reconstruction/h-dome transform. This has the advantage over simple thresholding that local extrema are more reliably detected, as they are then not only a function of absolute value, but also of local connectivity.

Our approach combines the strengths of these two works, as we found that combining Euclidean-distance-based heatmap regression with morphological reconstruction for seed extraction and subsequent watershed flooding to be very powerful. Furthermore, we buttress our pipeline with an emerging powerhouse of visual feature extraction: U-VixLSTM [28].

During training, we predict heatmaps generated by the Euclidean distance transform, with foreground values rescaled between -2.0 and +20.0 per cell and background set at -5.0. We use mean-squared error loss between predictions and targets. Furthermore, we apply reflective padding to both the training images and their corresponding heatmap labels. During training, the loss is computed only for pixels within this region, excluding padded areas. This strategy improves segmentation performance near image boundaries.

We also employ a specific sampling strategy, as we observe that random sampling without regard to cell density or illumination variability biases the training distribution towards unrepresentative samples, a common shortcoming of many existing segmentation pipelines. To address this, we designed a dynamic crop sampling scheme that alternates between object-centered crops, random image locations, unusually bright background patches, and unusually dark foreground patches, enhancing generalizability and improving foreground-background separation. Unusually bright background patches are identified before training by locating local maxima in the background within a neighborhood of an image crop size, and vice versa for the idiosyncratic foreground patches. For validation, however, we retain object-centered crops to compute the loss, as this approach is efficient and performs reliably in practice, without requiring full-image inference.

After training is complete, we perform tiled inference on validation images and optimize five watershed postprocessing hyperparameters using Optuna [45]: cell prominence (h-value for hdome transform [26]), minimum cell confidence, background threshold, preliminary Gaussian smoothing, and simple thresholding. Cell prominence defines the minimum value required for a cell to survive morphological reconstruction, which involves repeatedly dilating a marker image (created by subtracting the prominence value from the original image) while constrained by the original image as a mask. This pixel-connectivity-based process eliminates background variations smaller than the prominence threshold while preserving image peaks, since we threshold the output of morphological reconstruction with the cell prominence. Optional Gaussian smoothing may be applied before this step to improve heatmap cohesion. Then, watershed flooding [27] is applied to the inverted heatmap using h-dome-derived seeds and the background threshold to control label propagation. Finally, we threshold each cell obtained via this pipeline against a confidence threshold c. This eliminates background noise and can prevent cells from “leaking” into other parts of the image.

While morphological reconstruction is our primary choice due to its superior performance consistency across datasets, the h-dome transform can be substituted by simple thresholding for seed detection. This substitution enables a fully end-to-end, GPU-based inference pipeline, leveraging the embarrassingly parallel nature of the rainfall watershed implementation. However, because simple thresholding introduces minor performance fluctuations between different datasets, we treat this GPU-optimized pipeline as an auxiliary result; a comprehensive ablation study comparing performance with and without morphological reconstruction is provided in the Supplementary Material.

### Data preprocessing

We generally follow the preprocessing strategy described in [12]. In addition, we incorporate our own dataset of seven labeled image cubes from zebrafish neurons (N = 1,543) imaged with confocal microscopy, which pose particular challenges due to their low SNR and dense cell packing. All large images from which object crops are sampled are clipped between the 1st and 99.9th percentiles, then normalized to the [0, 1] range. We use the following bicubic upsampling factors and image crop sizes for the datasets Parhyale, Platy-CBG, Mouse Skull, and Zebrafish confocal, respectively: [7, 1, 1]: [128, 128, 128], [5, 1, 1]: [128, 128, 128], [1, 1, 1]: [128, 128, 128], [1, 1, 1]: [64, 64, 64]. We selected our crop sizes and upscaling factors to balance two practical needs: meeting the U-vixLSTM’s 32-pixel divisibility rule and ensuring the accuracy of the distance transform (EDT) labels through integer scaling. Upsampling factors in z were matched to the nearest integer of the anisotropy factor for each dataset, for example, 7 for Parhyale and 5 for Platy-CBG.

### Adaptive Optical Preprocessing

We prepend a Global Zernike Layer to the architecture to handle the varied image physics and point spread functions (PSFs) inherent in different microscopy setups. This layer models optical aberrations using Zernike polynomials [39, 40]. To adapt these traditionally 2D series for volumetric data, we apply Fischer decomposition to factorize polynomials on the unit ball into their radial and harmonic components [41]. Operating in the frequency domain, the layer applies a learned phase mask Φ to the Fourier-transformed input volume:

Xaberrated=F(x)⋅eiΦ

The mask Φ is a weighted sum of Zernike polynomials Zj. We found that optimizing only the coefficients for indices j∈{3,4,12}, representing longitudinal shift, defocus, and primary spherical aberration, produced the most consistent improvements in segmentation accuracy. To our knowledge, this is the first instance of using a learnable Zernike phase mask as a modular block within a deep imaging pipeline (Figure 4).

**Figure 3:**
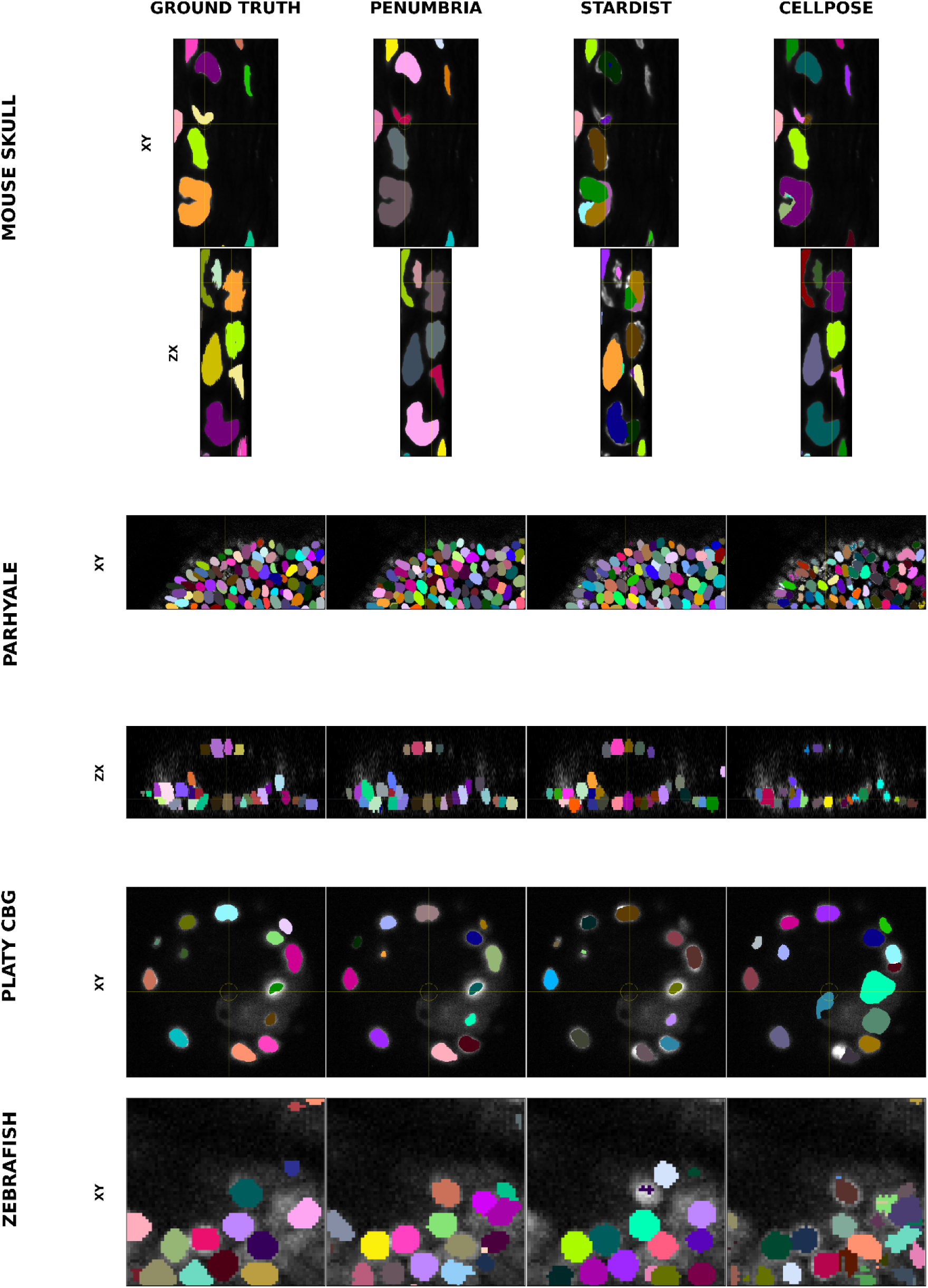
Visual comparison of the segmentation output of each method (Cellpose-SAM, Stardist-3D, and Penumbria) vs the ground truth on different datasets. Each color stands for a unique cell ID. Parhyale and Mouse Skull include an axial view for better inspection. Please note that cells or labels occasionally emerge in the next plane.

**Figure 4:**
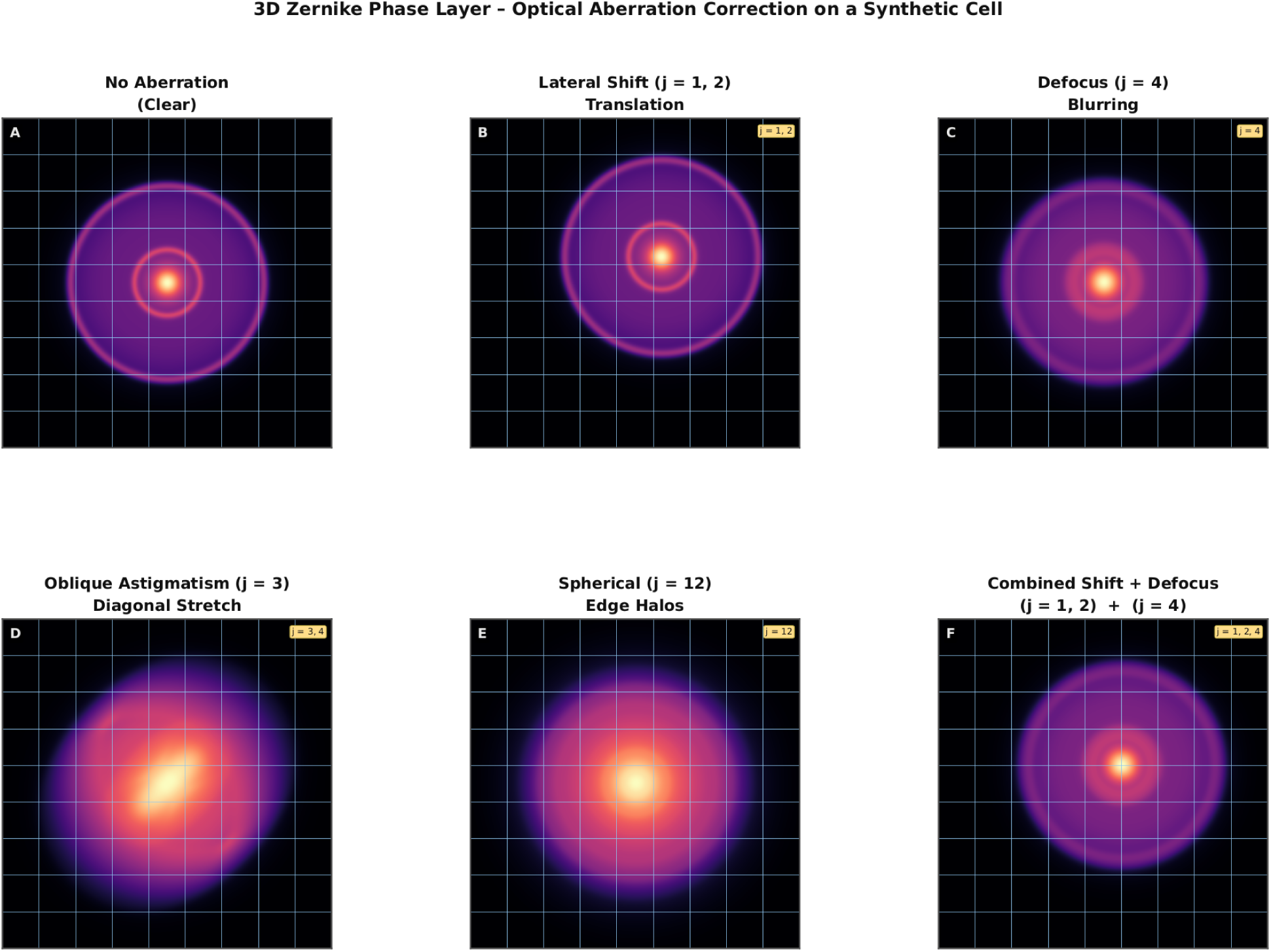
An illustration of the 3D Zernike phase layer that corrects optical aberrations caused by the microscope lens. For clarity, we limit our illustration to two dimensions. Before the signal reaches the main network, the layer learns to apply a phase correction on the unit ball, built from a weighted combination of Zernike polynomials. In this example, we demonstrate the effect of individual and summed Zernike coefficients on a synthetic cell.

**Figure 5:**
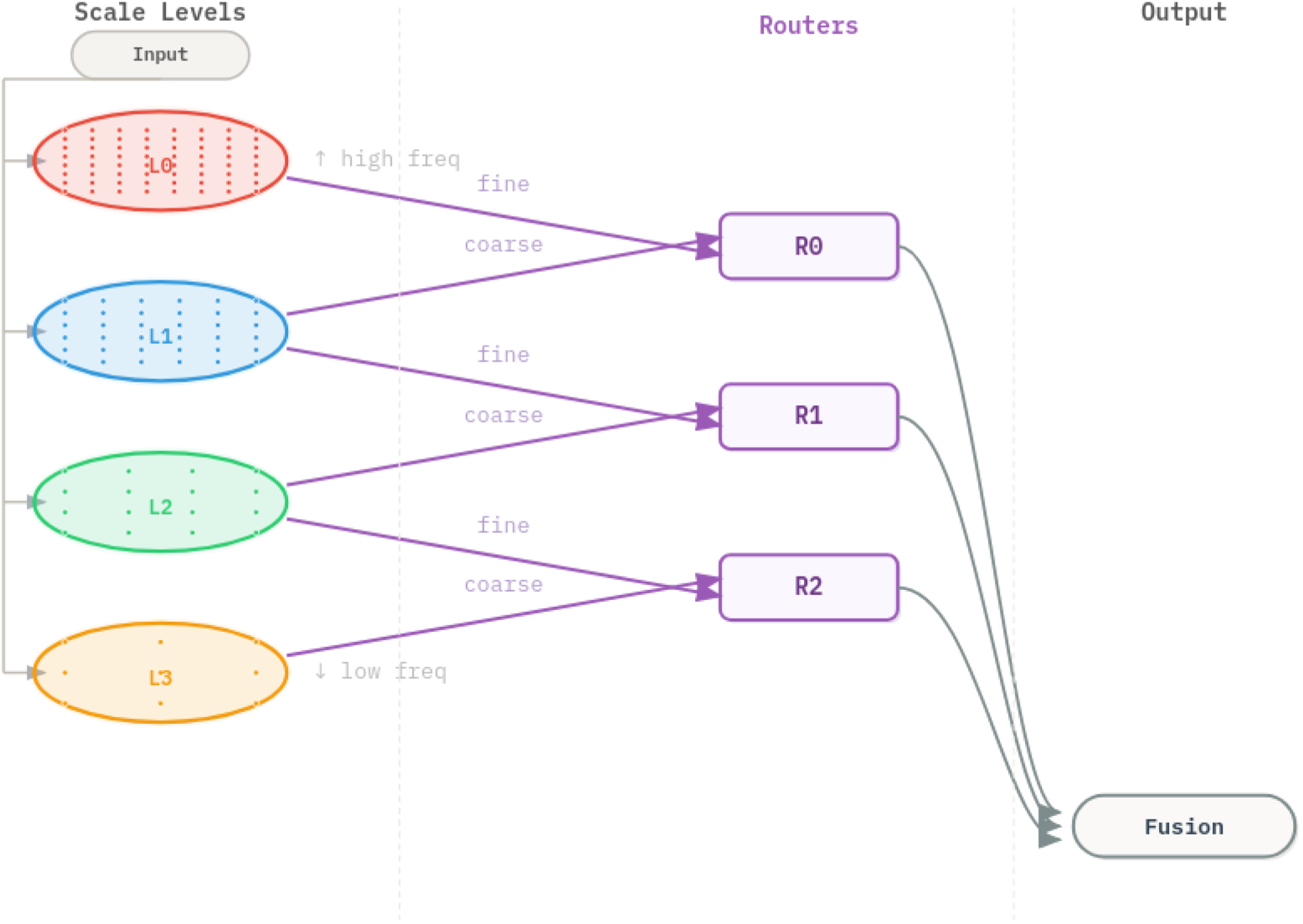
Illustration of the Scaled GeoCaps Layer. Taking inspiration from Capsule Networks, this layer seeks to find agreements not between part-whole relationships within an object, but different spatial scales represented by different sampling frequencies, here illustrated with the point density per module. Routers iteratively weave small-scale information into coarser visual features. Subsequently, feature maps are projected to the lowest resolution and fused in a linear layer.

We regularize the frequency response via Uniform Spectral Dropout. During training, we randomly mask individual frequency components with probability p. The DC component is always preserved to maintain stable global intensity scales, following established regularization techniques [46, 47].

Ablation results against the baseline network (Table 3) confirm the utility of the Zernike phase layer. Numerical gains remain robust, ranging from +0.11% in zebrafish neurons to +1.45% in Parhyale. The layer also generally stabilizes performance across data folds, except for the Mouse Skull dataset. As discussed in the following section, the Scaled GeoCaps (SGG) layer specifically addresses this outlier. These results were achieved by focusing on the fundamental optical properties defined by the j∈{3,4,12} Zernike moments. Note that the ablation was carried out against vanilla U-vixLSTM, while the SGG layer was ablated with the inclusion of the Zernike layer. This reflects how we developed these modifications in turn.

**Table 3:** Evaluation of stability and performance of the Zernike phase layer with moments j∈{3,4,12} compared to the baseline architecture. The average MaP boost is calculated over all IoU thresholds (0.1 - 0.9).

| Dataset | Overall Lead (Avg mAP) | Avg mAP Boost (%) | More Stable (Lower Std) | Std (Zernike) | Std (Base) |
| --- | --- | --- | --- | --- | --- |
| platy_cbg | **Zernike** | **+0.67%** | Zernike | 0.0548 | 0.0588 |
| parhyale | **Zernike** | **+1.45%** | Zernike | 0.0268 | 0.0325 |
| mouse_skull | **Zernike** | **+0.43%** | Base | 0.0865 | 0.0147 |
| zebrafish | **Zernike** | **+0.11%** | Zernike | 0.0330 | 0.0477 |

### Scaled Grid GeoCaps Layer

The Scaled Grid Geocaps (SGG) layer provides the xLSTM bottleneck with high-fidelity spatial information that standard encoder paths often lose. Instead of traditional feature aggregation, we adopt the routing mechanism from Capsule Networks [42]. While the original formulation focuses on semantic relationships between object parts, we repurpose it for geometric concordance by seeking agreement between identical spatial locations across different frequencies. A detection is only confident if evidence remains consistent across multiple scales simultaneously. This forced agreement promotes global coherence in the final heatmap output, effectively reducing split errors in large or elongated structures.

To provide the diverse perspectives required for this concordance, we use a bank of parallel convolutions to sample features at fixed grid locations. These scales use increasing strides for coarser grid sampling and increasing dilation rates to expand the effective receptive field. This design specifically addresses cases where cells are larger than the image crops fed into the network, such as in the Mouse Skull dataset, where the model requires a broad spatial context to maintain structural integrity.

To stabilize these representations and prevent performance drops on specific morphologies such as the Parhyale dataset, we also apply localized feature distillation. By providing sparse shape hints via a shallow MLP to random cells during 20% of training, we leverage the spatially anchored nature of the SGG layer to enforce bottleneck consistency. While we do not provide a formal improvement table for this tuning, we found that the SGG layer, in combination with this regularization, led to a 15% performance increase on the Mouse Skull dataset, while results for all other datasets remained within a noise margin of 0.5%.

### Core Architecture

Penumbria uses U-VixLSTM [28] as its core architecture, which features a multi-stage Unet design with VisionLSTM at the bottleneck, combining the strengths of both more local feature extractors (CNNs) and global view modelers (xLSTM). In particular, the network generates patch embeddings with various convolutional kernel sizes, followed by visionLSTM blocks to enhance the architecture’s global perception, thereby reducing false positives. This is combined with lateral UNet-style connections that concatenate the tensors from the previous decoder layer with appropriately shaped features from the encoder. The implementation of VisionLSTM for segmentation by [28] has so far received modest attention; however, both the results reported in their paper and our own experiments confirm that it consistently yields a measurable improvement across datasets compared to, for instance, vanilla Unet-based approaches or even Segformer3d [33]. Furthermore, it is not confined to Linux-based environments such as Segmamba [32], an important consideration given that biologists often work with Windows machines. Its only limitation is that it only allows isotropically shaped image volumes that are divisible by 32, but this can easily be mitigated by appropriate image resampling strategies, usually along the z dimension. We largely retain this architecture but augment it by replacing BatchNorm with Filter Response Normalization (FRN), and adding the Zernike phase layer on top of a Scaled GeoCaps (SGG) layer. We observe that FRN significantly reduces false positives, especially on datasets where brightness is the primary indicator of cell presence. It represents the optimal normalization choice because it preserves absolute values (unlike InstanceNorm), does not become unstable at smaller batch sizes (unlike BatchNorm, which is crucial for 3D segmentation), and has smaller memory requirements than GroupNorm [48, 49]. The Zernike phase layer yields modest yet stable improvements under different imaging conditions, and the SGG layer is particularly crucial for contexts where cells are too large to fit within the network’s effective receptive field.

### Training and Inference

We train the network using the SGD optimizer with a learning rate of 1e-4 and a momentum of 0.9. We explicitly hold out the same folds for validation and testing for Penumbria and the methods we benchmark against. Each algorithm is run 3 times on different folds for each dataset. We use a batch size of one to ensure consistency across datasets and mitigate out-of-memory errors. Training proceeds for up to 120,000 iterations with early stopping, and postprocessing parameters are optimized using Optuna [45] on the validation images for 300 iterations.

During inference, we use a sliding-window approach with overlapping tiles to mitigate boundary effects. Furthermore, we employ Euclidean-distance-weighted feathering to prevent intensity jumps and ensure a seamless transition between tiles. A sliding-window step size of half the predicted cube works best. Images are padded to a shape that is divisible by this tiling, using reflective padding to enhance realism. We use eight-fold test-time augmentation (TTA) in 3D by rotating the image crop axially and radially and averaging predictions. To save time, TTA is forgone on clear background patches (i.e., when all voxels belong to the darkest percentile of the neural output range). Before performing Optuna tuning on morphological postprocessing parameters for watershed seeding, the predicted heatmaps are resampled to their original resolution using bicubic interpolation. We trained Penumbria with mixed precision on both the MPCDF Raven cluster (A100s) and a local RTX 4090 (GPU memory requirements depend on configuration and dataset) to ensure it performs as well on a lab desktop as on an HPC.

### Use of AI tools

Large language models were used to assist with language editing and improving the clarity and structure of the manuscript. In addition, AI-based code generation tools were used to support the implementation of parts of the software. All generated content was reviewed and validated by the authors.

## Code availability

Our method is implemented in Python using PyTorch. Code is available at https://github.com/postnubilaphoebus/Penumbria under a non-commercial license.

## Acknowledgements

We thank Francesco Mistri for rerunning the CellPose algorithm on some datasets and Qing Wang for assistance with proofreading. We acknowledge support from the Max Planck Society.

## Author Contributions

L.S. conceived the original ideas, designed and implemented all modifications to the neural architecture, performed all experiments and analyses, and wrote the initial draft of the manuscript. J.D. provided regular supervision throughout the project, including extensive discussions on microscopy modalities, algorithmic trade-offs, and biological priors specific to the specimens studied; H.B. provided supervision, scientific guidance, and critical feedback, and secured funding and resources for the project.

## Notes

### Competing Interest Statement

The authors and/or their institution are in the process of filing intellectual property protection related to the technology described in this manuscript, including potential commercial applications. This represents a potential competing interest.

